# Oxytocin receptors influence the development and maintenance of social behavior in zebrafish (*Danio rerio*)

**DOI:** 10.1101/2021.06.26.449566

**Authors:** Anja Gemmer, Kristina Mirkes, Lukas Anneser, Tim Eilers, Caroline Kibat, Ajay Mathuru, Soojin Ryu, Erin Schuman

## Abstract

Zebrafish are highly social teleost fish and an excellent model to study social behavior. The neuropeptide Oxytocin is associated different social behaviors as well as disorders resulting in social impairment like autism spectrum disorder. However, how Oxytocin receptor signaling affects the development and expression kinetics of social behavior is not known. In this study we investigated the role of the two oxytocin receptors, Oxtr and Oxtrl, in the development and maintenance of social preference and shoaling behavior in 2- to 8-week-old zebrafish. Using CRISPR/Cas9 mediated *oxtr* and *oxtrl* knock-out fish, we found that the development of social preference is accelerated if one of the Oxytocin receptors is knocked-out and that the knock-out fish reach significantly higher levels of social preference. Moreover, *oxtr* ^-/-^ fish showed impairments in the maintenance of social preference. Social isolation prior to testing led to impaired maintenance of social preference in both wild-type and *oxtr* and *oxtrl* knock-out fish. Knocking-out one of the Oxytocin receptors also led to increased group spacing and reduced polarization in a 20-fish shoal at 8 weeks post fertilization, but not at 4. These results show that the development and maintenance of social behavior is influenced by the Oxytocin receptors and that the effects are not just pro- or antisocial, but dependent on both the age and social context of the fish.

## Introduction

Many species, including humans, live in groups to enhance their fitness – their lifetime reproductive success. Living in a social context offers many benefits like improved predator and food detection^1^, availability of mating partners, reduction of energy consumption^2^ as well as the opportunity to learn vital behaviors from conspecifics^3^. In order to optimize cohabitation within a group, different forms of social behavior evolved.

The zebrafish (*Danio rerio*), a small teleost fish, is a powerful animal model used in biomedical research, including drug discovery^4^, developmental biology^5,6^ and neurobiology^7,8^. Furthermore, zebrafish exhibit a variety of behaviors including avoidance^9^, foraging and hunting^10^, responses to stress^11^ and different forms of sociality^12-16^. Examples of sociality include mating behavior, aggressive behavior and other simpler behaviors that occur in groups. Zebrafish prefer to swim in cohesive shoals, a tendency that develops within the first weeks of age^17^. Swimming in close proximity to conspecifics, also called social preference, starts to develop as early as 1 - 2 weeks-post-fertilization (wpf)^18,19^. Although the development of shoaling behavior has been correlated with age-dependent changes in the dopaminergic and serotonergic system^20^, the mechanisms underlying the development and maintenance of social preference and shoaling behavior are not well understood.

The nonapeptide Oxytocin is a highly conserved neuropeptide, present in humans and with only minor alterations in most other animals^21^. The zebrafish orthologue Isotocin (abbreviated Oxt) differs by only two amino acids from human Oxytocin^21^. Next to its role in parturition^22^ and lactation^23^, Oxytocin has been described in the context of memory consolidation^24^ and nocifensive behavior^25^. Moreover, the association between Oxytocin and social behavior has been demonstrated in multiple studies^26-28^ and the human Oxytocin receptor is connected to neurodevelopmental disorders that are associated with impaired social behavior like autism spectrum disorder^29,30^. In mice, the Oxytocin receptor is expressed in different brain regions, e.g. in the hippocampus, amygdala, suprachiasmatic nucleus and prelimbic cortex^31^. As teleost fish like zebrafish experienced an additional round of whole genome duplication approximately 320-350 million years ago^32^, they possess two orthologous Oxytocin receptors, the Oxytocin receptor (Oxtr) and the Oxytocin receptor like (Oxtrl).

Making use of CRISPR/Cas9 generated *oxtr* and *oxtrl* knock-out lines, the primary objective of this study was to investigate the role of the Oxytocin receptors in the development and maintenance of social behavior, specifically social preference and shoaling behavior in zebrafish (*Danio rerio*).

## Results

To investigate the role of the Oxytocin receptors in development and maintenance of social behavior we compared the *oxtr* ^-/-^ and *oxtrl* ^-/-^ fish and their wild-type controls in two behaviors: social preference (assesses the fish’s preference for a social versus a non-social area) and shoaling.

### Social preference

To measure social preference, individual zebrafish larvae at different developmental stages were placed in a rectangular tank with one area where conspecifics could be seen through a transparent glass wall (Fig. 1a). Following a period of habituation, two conspecifics were added to the stimulus area of the tank and the behavior of the experimental fish was observed. We measured the time spent in proximity to conspecifics (social region-of-interest) as an indicator of a social preference for social preference. In wild-type zebrafish, this preference developed incrementally over the first 4 wpf and persisted up to 8 wpf (Fig. 1b,c). The *oxtr* ^-/-^ fish, by contrast, exhibited an early peak of maximal social behavior at 3 wpf. In contrast to the wild-type fish, this high sociability was not maintained but rather significantly (p = 1.65 × 10^−2^) decremented over 4-8 wpf (Fig. 1b,c). The development of social preference was also significantly (p = 3.49 × 10^−5^) altered in the *oxtrl* ^-/-^ fish, which also exhibited an early onset of social preference, reaching a peak precociously at 3 wpf (Fig. 1b,c). Furthermore, the maximal level of social preference exhibited by both Oxytocin receptor knock-out fish lines was substantially greater than the maximal level exhibited by the wild-type fish.

**Figure 1:**
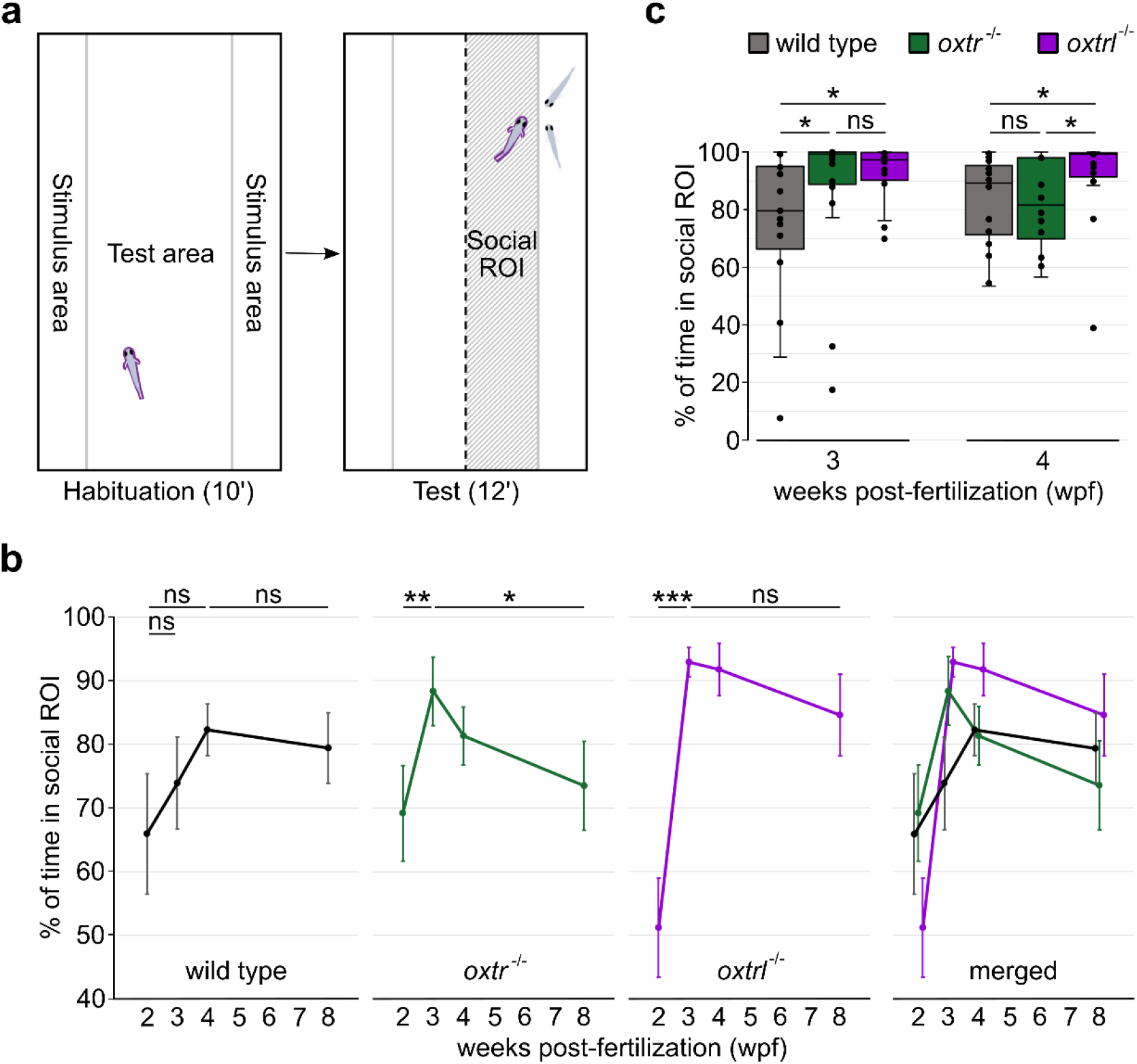
Oxytocin receptors affect the development, intensity and maintenance of social preference. **(a)** The behavioral chamber for social preference tests was composed of one test area and two stimulus areas, divided by transparent walls. After a habituation period, two stimulus fish were added to one stimulus area and the half next to the stimulus fish was defined as “social ROI” during analysis. **(b)** Social preference developed faster in both the *oxtr* ^-/-^ and *oxtrl* ^-/-^ fish and in *oxtr* ^-/-^ fish the maintenance of social preference was impaired. Values are reported as mean ± standard error of the mean (s.e.m.). **(c)** At 3 and 4 wpf, the level of social preference was increased in the *oxtrl* ^-/-^ fish. Fish of the *oxtr* ^-/-^ line also showed enhanced social preference at 3 wpf, but did not maintain this high level at 4 wpf. Number of replicates (n) can be found in Supplementary Table S1 and significance values in Supplementary Table S2. Significance is reported as *p<0.05, **p<0.01, ***p<0.001.

Isolation has been reported to alter social behavior in different species^33-36^. Therefore we next examined the effects of rearing in isolation on the development and expression of social preference in both wild-type and *oxtr* ^-/-^ and *oxtrl* ^-/-^ fish. We individually raised (in the absence of conspecifics) wild-type, *oxtr* ^-/-^ and *oxtrl* ^-/-^ fish from 2 days post-fertilization (dpf) to the day of experiment (Fig. 2a). In wild-type fish isolation rearing did not significantly (p = 2.03 × 10^−1^) affect the onset kinetics of social preference, but it dramatically diminished the maintenance of social preference typically observed at 8 wpf (Fig. 2b,c). Indeed, at 8 wpf isolation-reared wild-type fish failed to exhibit any significant (p = 3.26 × 10^−1^) social preference. Isolation rearing had a similar effect on the *oxtr* ^-/-^ fish: isolation did not alter the onset kinetics of social preference but led to an accelerated decline in social preference measured at 8 wpf. Interestingly, the *oxtrl* ^-/-^ fish showed the same pattern, but the accelerated decline in social preference following isolation rearing was precociously evident at 4 wpf (Fig. 2b,c). These data show that the social preference develops following isolation rearing in wild-type, *oxtr* ^-/-^ and *oxtrl* ^-/-^ fish, but this social preference is not maintained. The isolation rearing induced decline in social preference is evident earlier in *oxtr* ^-/-^ and *oxtrl* ^-/-^ fish.

**Figure 2:**
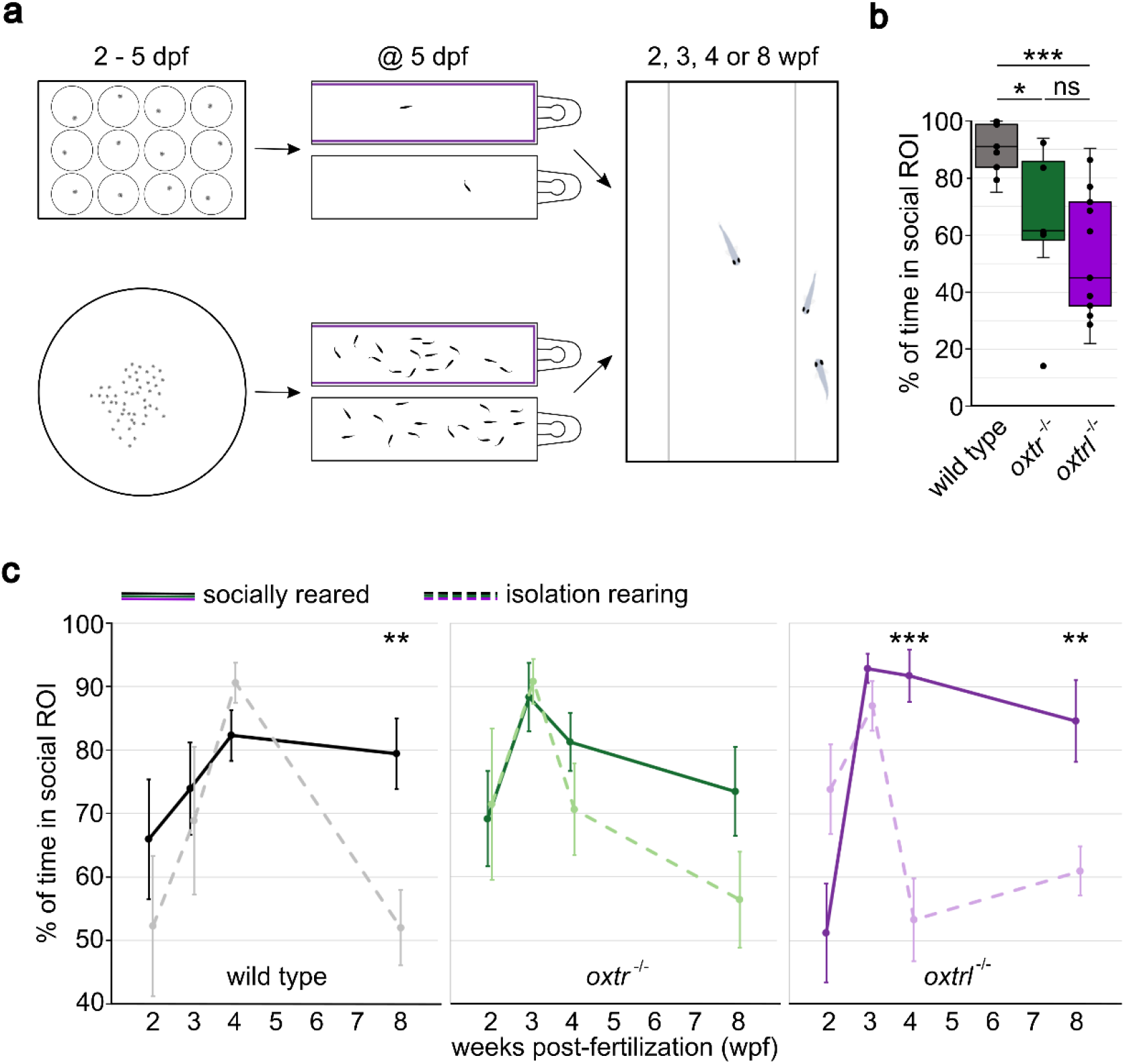
Isolation rearing impairs the maintenance, but not the development of social preference. **(a)** Rearing conditions: Experimental fish were raised in isolation (2-5 days post-fertilization (dpf) in 12-well plates, from 5 dpf until experiment in 1.1L tanks with visual barriers (here shown in purple) in every second tank) or with conspecifics (2-5 dpf in a dish with 145 mm diameter, from 5 dpf until experiment in 1.1L tanks with visual barriers in every second tank at densities of 10-15 fish/L). The social preference test was performed at 2, 3, 4 or 8 wpf. **(b)** *oxtr* ^-/-^ and *oxtrl* ^-/-^ fish showed decreased social preference at 4 wpf after isolation rearing. **(c)** The development of social preference was not influenced by isolation rearing. Similar to the phenotype observed for socially reared fish, knocking-out one of the Oxytocin receptors led to an accelerated development of social preference in isolated fish. Isolation rearing impaired the maintenance of social preference, but 4 wpf old *oxtr* ^-/-^ fish were less susceptible to isolation. Values are reported as mean ± standard error of the mean (s.e.m.). Number of replicates (n) can be found in Supplementary Table S1 and significance values in Supplementary Table S2. Significance is reported as *p<0.05, **p<0.01, ***p<0.001.

### Shoaling

We also investigated another form of social behavior: shoaling. To measure shoaling we used a setup which allowed 20 zebrafish to move freely in a round tank (see Supplementary Fig. S2a) for 30 minutes. We tested socially reared wild-type, *oxtr* ^-/-^ and *oxtrl* ^-/-^ fish at ages of 4 and 8 wpf (younger fish could not be reliably tracked owing to their small size). To quantify shoaling, we analyzed the nearest-neighbor distance, inter-individual distance and farthest-neighbor distance of each fish as well as the cumulative shoal distance and the polarization parameter obtained from principal component analyses of individual videos (see Materials and methods). At 4 wpf, neither of the Oxytocin receptor knock-outs exhibited shoaling characteristics that were significantly different from their associated wild-type controls (p(*oxtr* ^+/+^ vs. *oxtr* ^-/-^) = 4.74 × 10^− 1^ (a), 3.85 × 10^−1^ (b), 4.74 × 10^−1^ (c) and 4.44 × 10^−1^ (d), p(*oxtrl* ^+/+^ vs. *oxtrl* ^-/-^) = 2.11 × 10^−1^ (a), 8.64 × 10^−2^ (b), 7.86 × 10^−2^ (c) and 5.28 × 10^−2^ (d)) (Fig. 3a-d). At 8 wpf, however, both knock-out lines exhibited significant differences in their shoaling features, all consistent with the general phenotype of a less cohesive shoal (p(*oxtr* ^+/+^ vs. *oxtr* ^-/-^) = 5.48 × 10^−4^ (a), 4.59 × 10^−3^ (b), 8.19 × 10^− 3^ (c) and 5.24 × 10^−3^ (d), p(*oxtrl* ^+/+^ vs. *oxtrl* ^-/-^) = 2.75 × 10^−2^ (a), 1.34 × 10^−2^ (b), 1.27 × 10^−2^ (c) and 1.79 × 10^−2^ (d)) (Fig. 3a-d). Taken together, these two experiments revealed that both Oxytocin receptors play a crucial role in the development of social preference and shoal cohesion.

**Figure 3:**
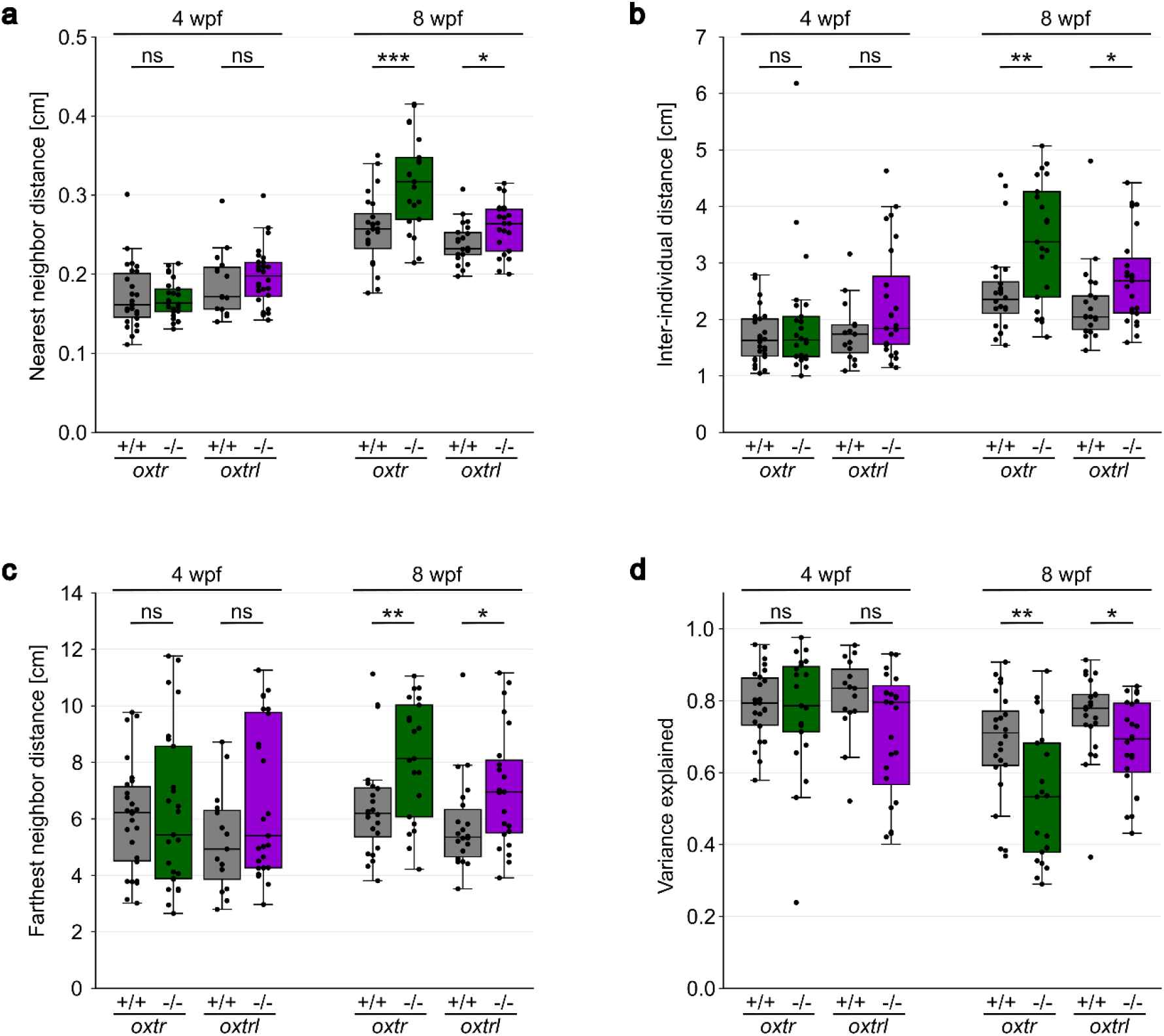
Shoal cohesion and polarization is positively influenced by the Oxytocin receptors at 8 wpf. **(a)** The absence of either Oxytocin receptor led to increased nearest neighbor distance at 8 wpf, but not at 4 wpf. **(b)** The inter-individual distance was enlarged in both Oxytocin receptor KO lines at 8 wpf, but not at 4 wpf. **(c)** *oxtr* ^-/-^ and *oxtrl* ^-/-^ showed increased farthest neighbor distance at 8 wpf, but not at 4 wpf. **(d)** Coordinated swimming, represented by the variance explained, was reduced in both knockout groups at 8 wpf but remained unaffected at 4 wpf. Number of replicates (n) can be found in Supplementary Table S1 and significance values in Supplementary Table S2. Significance is reported as *p<0.05, **p<0.01, ***p<0.001.

## Discussion

Many studies^37-39^ have demonstrated a connection between Oxytocin and social behavior, but the exact role of the two Oxytocin receptors in development and maintenance of social behavior remained to be clarified. Our data revealed that preference for companions develops gradually in wild-type zebrafish in the first few weeks, reaching a maximum social preference at 4 wpf, which is then maintained at a high level until at least 8 wpf. If one of the Oxytocin receptors (Oxtr or Oxtrl) was deleted, the development of social preference was accelerated, reaching its maximal level precociously at 3 wpf. Although Autism spectrum disorder is caused by multiple factors, mutations in the Fragile X mental retardation 1 (*FMR1*) gene^40^ as well as specific single-nucleotide polymorphisms (SNPs) of the human *OXTR*^*29*^ gene have been reported to cause autism-like symptoms. We have shown here that the development of social preference is affected by Oxytocin receptors in a distantly-related vertebrate, the zebrafish. It is interesting to note, that a recent study reported that the development of social preference is also sped up in *fmr1*^-/-^ zebrafish^41^.

Isolation rearing of zebrafish did not affect the time course of the development of social preference, but significantly affected its maintenance. The isolation-induced decline in social preference was accelerated in both of the Oxytocin receptor knock-outs. RT-PCR of *oxt* expression in socially- and isolation-reared fish revealed that isolation led to significantly reduced *oxt* levels at 8 wpf, but not at 4 wpf (see Supplementary Fig. S1c). Furthermore, and in contrast to socially-reared fish, isolation-reared fish did not show significantly higher *oxt* expression with increasing age. These data are consistent with the observed behavioral changes after isolation rearing. Another possible explanation for the dramatically reduced social preference level at 8 wpf might be the downregulation of *pth2*, as isolation leads to reduced expression of this neuropeptide^42^.

Our results differ from the work of Landin and co-workers, who tested the social preference of adult and 3-week-old zebrafish after application of the Oxytocin receptor antagonist L-368,899, a specific inhibitor of both Oxtr and Oxtrl^43^. At both developmental stages they detected a decreased social preference. However, the social preference chamber they used did not allow the simultaneous visual access of both stimulus areas. In this chamber the fish have to pass through a small corridor to move between the two compartments which may provoke an anxiety-like state. As Oxytocin has an anxiolytic effect^44^ it is possible that the Oxytocin receptor antagonist leads to increased anxiety. Unfortunately, Landin and colleagues did not test anxiety-like behavior of larval zebrafish. Moreover, as the antagonist was added to the swim water, it is difficult to predict the concentration within the animal and incomplete inhibition of the receptors cannot be ruled out.

Unlike the Landin et al. study, a previous study by Zimmermann and colleagues did not find a significant decrease in social preference after intraperitoneal injection of L-368,899. The conflicting results in the two studies were attributed to the use of different concentrations of the antagonist, 0.01ng/g^45^ and 100µg/g^43^. In contrast to these two studies, we used CRISPR/Cas9 generated genetic mutants lacking only one of the two Oxytocin receptors allowing us to dissect their distinct individual roles throughout development. Both receptors influenced the development of social preference, but in contrast to *oxtrl* ^-/-^, *oxtr* ^-/-^ fish did not maintain a high level of social preference as they became older. Wircer and colleagues^46^ described a subpopulation of oxytocinergic neurons, which are located in the posterior tuberculum and express *oxtr*. Ablation of these neurons led to reduced social preference in adult zebrafish^46^. Interestingly, a study by Ribeiro and colleagues^47^ showed impaired social recognition but unchanged social preference in a single Oxytocin receptor (*oxtr*^wz16/wz16^) knock-out zebrafish line. Difference in the results obtained between Ribeiro et al. and our study may be due to the difference in experimental condition or age as they used a different behavioral chamber for the social preference test and three- to six-month old fish. *oxtrl* knock-out fish were not tested in their study. Different studies^48-51^ with *Oxtr* knock-out mice have shown decreased social behavior and impaired social recognition with enhanced aggression. Remarkably, the deficits in social behavior were also present in heterozygous *Oxtr* ^+/-^ mice^51^. In these studies, adult mice were tested. It is possible that the significant decrease in social preference of *oxtr* ^-/-^ fish continues until adulthood, resulting in impaired social preference of adult *oxtr* ^-/-^, but this remains to be tested with knock-out fish. In line with the Ribeiro et al. study^47^, a study with Oxytocin receptor knock-out prairie voles^52^ showed impaired social recognition in *Oxtr* ^-/-^, but did not find significant changes in social behavior. Our results further show that Oxtr and Oxtrl do not have same functions since *oxtr* ^-/-^ and *oxtrl* ^-/-^ knockouts exhibit different degrees of social preference throughout development. Importantly, when considering the role of oxytocin receptors, one may need to evaluate the potential role of Vasotocin, the zebrafish orthologue to mammalian Vasopressin, which can also bind to Oxtr and Oxtrl, though with a much lower affinity than Oxt^43^.

As *oxtrl* ^-/-^ fish show enhanced social preference level at 3 and 4 wpf, we expected them to shoal more tightly, with decreased group spacing compared to wild-type. The increase in social preference was not, however, accompanied by changes in the shoaling parameters of nearest-neighbor, inter-individual, farthest-neighbor distances and the polarization parameter “variance explained” in either the *oxtr* ^-/-^ and *oxtrl* ^-/-^ at 4 wpf. And yet, at 8 wpf the nearest-neighbor, inter-individual, and farthest-neighbor distances were significantly increased while the variance explained significantly decreased in both *oxtr* ^-/-^ and *oxtrl* ^-/-^. These data indicate less polarized shoaling behavior with increased group spacing in *oxtr* ^-/-^ and *oxtrl* ^-/-^. In the Landin et al. study, intraperitoneal injection of L-368,899 also led to enhanced nearest-neighbor, inter-individual, and farthest-neighbor distances in shoals of four adult fish^43^, supporting a pro-cohesive role of the two Oxytocin receptors in the organization of shoals with fish older than 8 wpf. The results from our shoaling experiments show a pro-cohesive and pro-polarization effect of Oxytocin via both Oxytocin receptors at 8 wpf, with a more pronounced effect of Oxtr, but this effect was not present at 4 wpf. As the shoaling behavior develops continuously until adulthood^17^, the most parsimonious explanation for both sets of results is that Oxytocin receptors are likely to be important for shoal organization at later stages of development, whereas other signaling pathways may be able to compensate at 4 wpf. Similar to the social preference, both Oxytocin receptors modulate shoaling behavior, but to different degrees.

In a study by Tang and colleagues^53^ the shoaling behavior of 90 CRISPR/Cas9 generated knock-out lines (for different genes) was compared (6 adult fish per shoal). This study revealed the influence of multiple genes on collective behavior of zebrafish – in particular on swimming speed, group spacing and polarization. Tang et al. further showed that a single gene does not necessarily affect swimming speed, group spacing and polarization, but can influence only one or two of these shoaling parameters. Our data are consistent with this finding as we did not detect significant changes in the cumulative shoal distance at 8 wpf (see Supplementary Fig. S2b), whereas the group spacing and polarization parameters were significantly altered at this age. At first glance, the observations that the Oxytocin receptor mutant fish develop a preference for social companions precociously, yet exhibit poor coordination and shoaling behavior at 8 wpf may seem inconsistent. However, this may be explained by the observations of a decline in preference for social companions over ontogeny. *oxtr* ^-/-^ mutants, for example, show a larger effect in diminished shoaling parameters at 8 wpf compared to *oxtrl* ^-/-^ which agrees with the larger reduction in the preference for social companions at this age. The modest or no change in shoaling parameters at 4 wpf could reflect either a technical limitation of this assay at an early age, or, that there may be a “ceiling effects” to detect improved shoaling characteristics.

Taken together, our results show that the Oxytocin receptors play an important role in the development and maintenance of zebrafish social behavior and the impact of Oxytocin signaling depends on the age and environment of the fish.

## Materials and methods

The materials and methods section follows the recommendations in the ARRIVE2.0 guidelines^54^.

### Study design, sample size and exclusion criteria

The number of biological replicates for each experiment can be found in Supplementary table S1.

For the investigation of social preference, we tested 18 fish per group. The sample size was determined prior to experiments using the E-equation method^55^. Data was excluded from further analysis if the test fish’s average swimming speed was below a threshold (79/411). This threshold was calculated for each age group by determining the average swimming speed of each genotype group (wild type, *oxtr* ^-/-^ or *oxtrl* ^-/-^) and multiplying the smallest with 0.6. The fish of three genotype groups (*oxtr* ^+/+^;*oxtrl* ^+/+^ = “wild type”, *oxtr* ^-/-^;*oxtrl* ^+/+^ = “*oxtr* ^-/-^“ and *oxtr* ^+/+^;*oxtrl* ^-/-^ = “*oxtrl* ^-/-^”) were reared either in isolation or with conspecifics at 10-15 fish/L density and were tested for their social preference at age of 2, 3, 4 or 8 wpf.

In the 30-minute long shoaling experiment, shoaling parameters were analyzed in a shoal of 20 fish at age of 4 or 8 wpf in three different genotypes (*oxtr* ^+/+^*oxtrl* ^+/+^ = “*oxtr* ^+/+^” or “*oxtrl* ^+/+^”, *oxtr* ^-/-^*oxtrl* ^+/+^ = “*oxtr* ^-/-^“and *oxtr* ^+/+^*oxtrl* ^-/-^ = “*oxtrl* ^-/-^”). In contrast to social preference, we detected a significant difference (variance explained(8 wpf): p(*oxtr* ^+/+^ vs. *oxtrl* ^+/+^) = 2.33 × 10^−2^, cumulative shoal distance(4 wpf): p(*oxtr* ^+/+^ vs. *oxtrl* ^+/+^) = 1.79 × 10^−2^) in some shoaling parameters between the homozygous wild-type cousins of *oxtr* ^-/-^ and the homozygous wild-type cousins of *oxtrl* ^-/-^. Therefore, we analyzed the wild-type zebrafish as two genotype groups, “*oxtr* ^+/+^” and “*oxtrl* ^+/+^”, respectively. We tested 23-28 socially reared (10-15 fish/L) shoals per group.

Test fish were used only once, sacrificed and genotyped after the experiment. If the fish/shoal was identified as heterozygous (16/611) it was excluded from further analysis.

### Experimental animals and generation of knock-out

The *oxtr* and *oxtrl* genes were mutated by Ajay Mathuru and Caroline Kibat using the sgRNA:Cas9 system described in ^**56**^. Two CRIPSR targets (<u>GGA</u>AGTTACCGTGTTGGCCT and <u>GGC</u>TGATAAGCTTTAAAATA for *oxtr*; GTGCGTCCTTGTGGCCATCC and GGGGGGATTTTGTTCAGCCC for *oxtrl*) for each gene were identified using ZiFit (http://zifit.partners.org/zifit/). Customized sgRNAs with 20 nucleotide sequence complementary to a target site were synthesized by first cloning the target sequences into the expression construct [pDR274](https://www.addgene.org/browse/article/6371/) (Addgene #42250) and then *in vitro* transcribed using T7 promoter according to the manufacturers protocol (Thermo Fisher # AMB13345). The Cas9 mRNA was transcribed from linearized plasmid [MLM3616](https://www.addgene.org/42251/) (Addgene #42251) following the manufacturers protocol (Thermo Fisher # [AMB1344](https://www.thermofisher.com/order/catalog/product/AM1344?SID=srch-srp-AM1344#/AM1344?SID=srch-srp-AM1344)). The sgRNA:Cas9 RNAs cocktail (containing 12.5ng/μL sgRNA and 300ng/μL Cas9) were injected into single-cell embryos of AB wild-type background. The efficiency of the CRISPR targets and quality of the Cas9 endonuclease were determine 24 hours post injection of sgRNA:Cas9 by PCR on 10% of the injected embryos. The remaining embryos were raised to adulthood and genotyped at three months post injection. A total of 64 individuals were genotyped. The sequence of the genotyping primer set is in detailed in the genotyping methods section. The PCR products were cloned into [pGEMT](https://www.promega.sg/products/pcr/pcr-cloning/pgem-t-easy-vector-systems/?catNum=A1360) (Promega #1360) and sequenced to verify the mutations. The F0 mutants were then then outcrossed to Danio Reds (https://doi.org/10.1016/S0006-291X(03)01282-8). F1 fish were in-crossed to establish germline transmitting homozygous mutant lines.

The lines are named *oxtr* ^ync02^ and *oxtrl* ^ync03^, in this paper abbreviated as *oxtr* ^-/-^ and *oxtrl* ^-/-^, respectively. The homozygous wild-type cousins of these fish were used as wild-type control. A compensation of the knocked-out receptor by its orthologue Oxytocin receptor was not detectable (see Supplementary Fig. S1b).

On the day of experiment the test fish were 2, 3, 4 or 8 wpf old, depending on the age group. We used experimental fish with a total body length comparable to the total body lengths described in the Zebrafish Book^57^: 6 mm (2 wpf), 8 mm (3 wpf), 10 mm (4 wpf) and 14 mm (8 wpf). In the social preference test the stimulus fish (two per test fish) had the same age and similar size than the test fish. As it is difficult (juveniles) to impossible (larvae) to distinguish males and females at the developmental stages used in this study, fish became experimental fish independent of their sex.

### Zebrafish housing and husbandry

Up to 5 dpf, the larvae were kept in a 28.5°C incubator with a 14/10 light-dark cycle. At 2 dpf, eggs were either individually isolated in a 12-well plate (3 mL E3 medium [5 mM NaCl, 17 mM KCl, 0.33 mM CaCl_2_, 0.33 mM MgSO_4_] per 22 mm diameter well) or kept in pertri dishes (145 mm diameter, filled with 150mL E3 medium) in groups of 50.

From 5 dpf on, they were raised isolated (including visual barriers placed in every second tank, see Fig. 2a, center) in 1.1L ZebTEC tanks or in groups of mixed sexes at densities of 10-15 fish/L in 1.1L ZebTEC tanks (social preference test) or 3.5L ZebTEC tanks (shoaling test).

From 5 dpf on, zebrafish were housed in a ZebTEC Active Blue Stand Alone System, with the water temperature of 28.5°C +/- 1°C, the pH = 7.4 +/- 0.3 and the conductance = 650 µS +/-100. The light-dark cycle in the fish facility is 14h-light/10h-dark with a 30 minute twilight phase. Fish were fed three times per day: Up to 10 dpf with vinegar eelworms (*Turbatrix aceti*, bred in house), CAVIAR 50-100 (SAFE) and crushed Gemma Micro 75 (Skretting), from 10 to 20 dpf with vinegar eelworms, brine shrimp (*Artemia salina, Ocean nutrition V154019*) and CAVIAR 50-100, from 21 dpf to 56 dpf with brine shrimp and CAVIAR 100-200 (SAFE).

### Experimental procedures

#### Social preference test

Behavioral chamber (height 10 mm) for testing social preference consisted of the test area (25 × 75 mm (for 2 to 4 wpf) and 50 × 75 mm (for 8 wpf)) and the stimulus areas (8 × 75 mm (for 2 to 4 wpf) and 16 × 75 mm (for 8 wpf)). The behavioral chamber was placed on a screen, providing white backlight illumination and a camera positioned above recorded the fish’s movements with 20 frames per second using the program “Pylon recorder”. To reduce visual and acoustic disturbances, the setup was surrounded by a black, sound-absorbing box (LBH: 450 × 670 × 850 mm). Prior to the experiment the test and stimulus fish were moved from their home tank to a 1L breeding tank (TECNIPLAST, Part Number: ZB10BTE) with a nursery insert (TECNIPLAST, Part Number: ZB300BTI) and kept in a 28.5°C incubator (test and stimulus fish in different tanks). After each test, the behavioral chamber was cleansed with hot water (∼60°C) and refilled with fresh ZebTEC Stand Alone system water (28.5°C). The fish were moved using a disposable 3ml pipette (the tip was cut off to generate a sufficient big diameter) by elevating the nursery insert and then transferred to the center of test area (test fish) or edge of stimulus area (stimulus fish) of the behavioral chamber. At 8 wpf the chamber was covered with the lid of a 145 mm dish to prevent fish from jumping out. The test fish habituated to the chamber without any other fish for ten minutes followed by a 12 minutes test phase in which two stimulus fish of same age and size were placed to one of the stimulus areas. A transparent wall between test and stimulus area allowed visual access to the stimulus fish whereas a white opaque wall determined the rear side of the stimulus area.

#### Shoaling experiment

The shoaling setup (see Supplementary Fig. S2a) consisted of a white round behavioral chamber with a diameter of 70 cm which was surrounded by 29.5°C water (heated by a pump) to prevent cooling of the water inside the behavioral chamber during the 30 minutes of experiment. A high-resolution camera (Basler acA4112-30um), positioned 73 cm over the chamber, recorded the fish with 30 frames per second using the program “Pylon recorder” and 1200 white LEDs in the ceiling provided the necessary illumination (approximately 650 Lux). A white box surrounding the setup reduced visual and acoustic disturbances of the fish. For each replicate the chamber was completely emptied, dried off and refilled with 2.4 L (4 wpf)/ 3.0 L (8 wpf) fresh 28.5°C warm Stand Alone system water. In the morning of an experimental day, the home tanks were moved from the ZebTEC stand alone to a 28.5°C incubator. Approximately 5 minutes before the experiment started, 20 fish of same age and similar size were transferred from their home tank and to a 1L breeding cage (TECNIPLAST) with a nursery insert (TECNIPLAST, Part Number: ZB300BTI). With the nursery insert all 20 fish of one shoal were moved simultaneously to the center of the behavioral chamber. The recording of shoaling behavior started immediately and continued for 30 minutes.

### Genotyping

In order to identify homozygous knockouts (KO) and homozygous wild types (wt) for generation of experimental fish, fin clip PCR was performed according to the ZIRC genotyping protocols (https://zebrafish.org/wiki/protocols/genotyping). Additionally, each experimental fish was genotyped after the experiment. The following primer sequences were used.

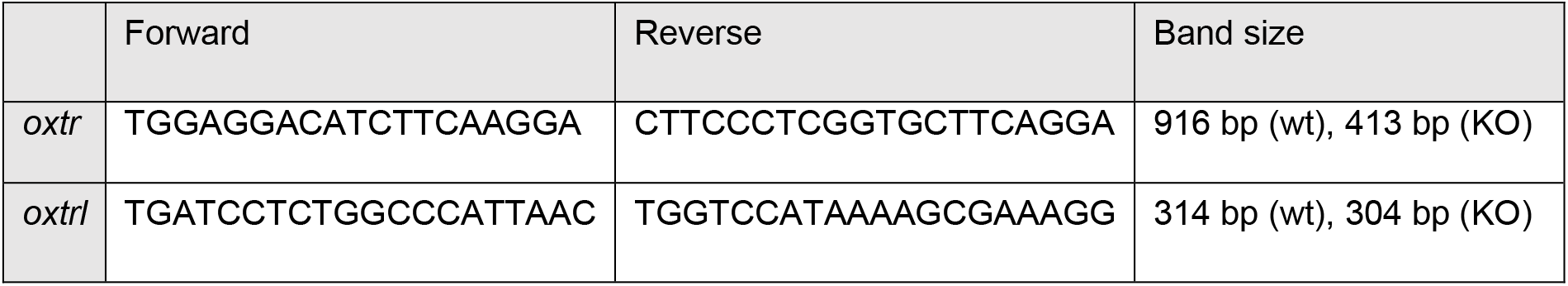

The PCR products were loaded on a 1% (*oxtr*) or 2% (*oxtrl*) agarose gel and run at 100V for 30 minutes (*oxtr*) or > 2 hours (*oxtrl*). But as the difference between *oxtrl* ^+/+^ and *oxtrl* ^-/-^ is only 10 base pairs, PCR products were also sequenced using the forward primer.

### Real-time PCR

RNA was extracted from isolated brains as described before^42^ and using the QuantiTect Reverse Transcription Kit (QIAGEN Cat. No. 205311) 200ng of RNA was transcribed into cDNA. 5μL of 1:10 diluted cDNA template were mixed with 1.3μL primers (10μM) and 6.25 μL SYBR Green PCR master mix (Applied Biosystems/Thermo Fisher Art. No. 4309155). The cycling parameters were 10 minutes at 95°C, followed by 40 cycles of denaturation (15 seconds at 95°C) and amplification (60 seconds at 60°C). A Real Time PCR System (Applied Biosystems/Thermo Fisher Art. No. 4376600) was used for qRT-PCR. The fluorescence threshold was set to 0.9 for all experiments and genes and we used TATA box binding protein (*tbp*) as reference gene. We used the ΔC_T_ method and the following primer sequences.

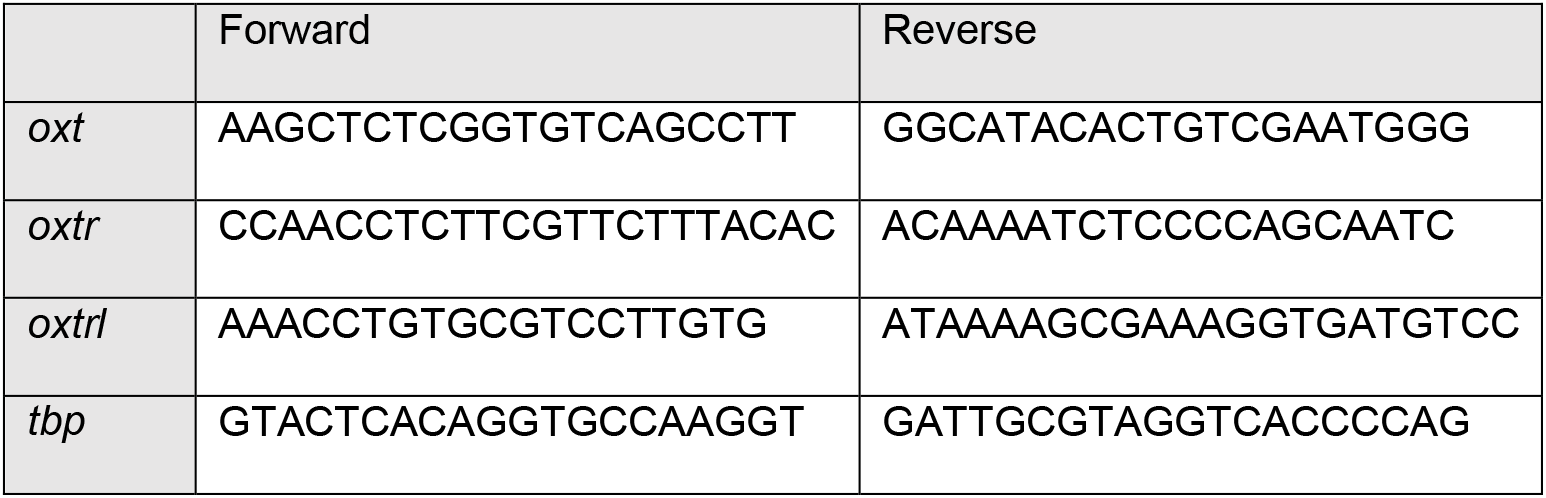

### Outcome measures

In the social preference test, the location of the test fish was measured in each frame of habituation and during the last 10 minutes of test, as the stimulus fish needed up to 2 minutes after being transferred to the stimulus area to exhibit normal social behavior. During analysis the test area was divided into two regions of interest (ROI). The “social ROI” was an area next to the stimulus fish whereas the “antisocial ROI” was on the opposite site of the test area. The percentage of total time was determined, in which the fish was located in the “social ROI”. The average swimming speed in absence (habituation) or presence (test) of stimulus fish was calculated and used as exclusion criteria for fish with reduced motion (e.g. because of freezing).

In the shoaling experiments all twenty fish of each shoal were tracked using TRex^58^ (tracking threshold 15 (4 wpf) and 50 (8 wpf)) and the nearest-neighbor distance, the inter-individual distance and the farthest-neighbor distance of each fish were calculated and cumulative shoal distance and variance explained as parameter for coordinated swimming were determined. For determination of the variance explained, x- and y-components of all individual trajectories (20 per shoal) were first assembled in a 40 × 54000 (30 minutes with 30 frames per second) matrix, whose covariance matrix was then eigen decomposed to obtain principal components (PCs). We found the first two PCs corresponded to the x- and y-component of the shoal centroid trajectory over time.

### Statistical methods

Significance is reported as follows: *p<0.05, **p<0.01, ***p<0.001.

All significance values can be found in Supplementary Table S2.

Normal distribution of data was checked using the Kolmogorov-Smirnoff-Test which revealed a non-normal distribution. Unpaired data was tested with a Kruskal-Wallis Test followed by a post-hoc Mann-Whitney-U Test (= Wilcoxon rank-sum test). The statistical analyses were carried out using python (from scipy.stats: kstest, kruskal and mannwhitneyu) or MATLAB (kstest, signrank, kruskalwallis and ranksum). In the social preference test n = 332 fish were included in the analysis (79 excluded), in the shoaling experiment n = 184 shoals were included in the analysis (16 excluded).

### Blinding and randomization

The investigators were not blinded to the genotype of the experimental fish, but both experiments were recorded with an overhead camera and analyzed using a tracking software (custom written tracking and analysis script for social preference and TRex^58^ plus a custom written python script for shoaling) to provide an unbiased data analysis. The test fish were chosen randomly from their home tank group and in the social preference test the location of stimulus fish was changed between replicates in a random order. The experiments were performed from 8:00 AM to 5:00 PM and the order of genotype groups per day was changed between experimental days to minimize a potential circadian confounder. Furthermore, the position of tank during rearing was chosen in a way that all replicates of one experimental group were reared in all possible heights/light conditions.

## Ethical statement

All procedures were conducted in accordance with the institutional guidelines of the Max Planck Society and were approved by the Regierungspräsidium Darmstadt, Germany (governmental ID: V 54-19 c20/15-F126/1013 and V54-19 c20/15-F126/1016).

## Data availability

The datasets generated and analyzed during this study and the custom written scripts are available from the corresponding author on request.

## Acknowledgements

We thank Anett-Yvonn Loos, Annette Hüttling, Dmitrji Burgard and Hans Siegert for the zebrafish husbandry and Fabian Bayer, Thomas Maurer, Guido Schmalbach, Andreas Umminger and Erik Papuschin for the manufacturing of our behavioral setups. We thank IMCB Zebrafish Facility staff for husbandry and Suresh Jesuthasan for sharing the *oxtr* and *oxtrl* knock-out lines with us. Furthermore we are grateful to Friedrich Kretschmer for the development of the program “Pylon Recorder” and Florian Vollrath for the tracking software and MATLAB script used for analysis of the social preference test. Finally we thank Helene Will for post-hoc genotyping of the experimental fish and Anett-Yvonn Loos for fruitful discussions. A.S.M. was supported by Yale-NUS College and Ministry of Education, Singapore through a Start-Up Grant.

## Contributions

A.G. and E.M.S. conceived the project. A.G., S.R. and E.M.S. designed the experiments, C.K. and A.S.M. generated the *oxtr* and *oxtrl* knock-out fish. A.G. and K.M. conducted the experiments, bred and genotyped the experimental fish, L.A. and T.E. wrote the python script for analysis of the shoaling experiments, A.G., L.A. and T.E. analyzed the data, A.G. and E.M.S. wrote the manuscript and prepared the figures. All authors reviewed the manuscript.

## Additional information

### Declaration of interests

The authors declare no competing interests.

**Supplementary Figure S1:**
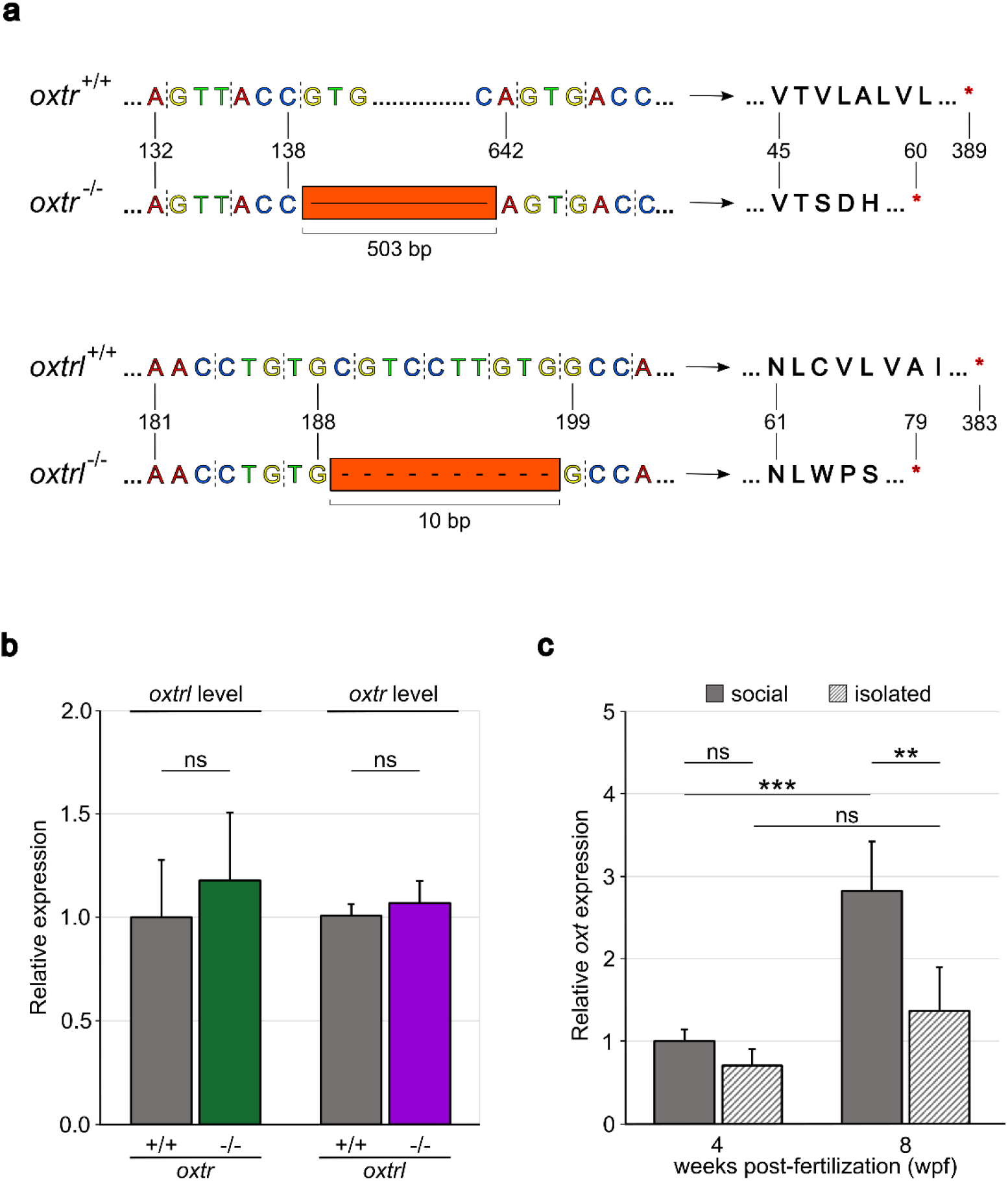
Details of the CRISPR/Cas9 generated KO, potential compensation and effect of isolation on *oxt* transcription. **(a)** A 503 base pair deletion in the *oxtr* gene and a 10 base pair deletion in the *oxtrl* gene caused frameshifts leading to premature stop codons. The resulting mutant Oxtr protein size is 59 amino acids (wild-type Oxtr protein is 388 amino acids) and mutant Oxtrl protein size is 78 amino acids (wild-type Oxtrl protein is 382 amino acids). From the different receptor parts (N- and C-terminus, seven transmembrane helices, three intracellular and three extracellular loops) only the N-terminus and parts of the first transmembrane helix were present in mutant Oxtr and Oxtrl, respectively. **(b)** The knocked-out receptor was not compensated by enhanced expression of the intact cognate receptor (Oxtrl in *oxtr* ^-/-^, Oxtr in *oxtrl* ^-/-^). **(c)** Isolation rearing led to significantly decreased *oxt* expression at 8 wpf. Under social conditions, wild-type zebrafish expressed Oxytocin at significantly higher levels with increasing age. This effect is significantly reduced after isolation rearing. Number of replicates (n) can be found in Supplementary Table S1 and significance values in Supplementary Table S2. Significance is reported as *p<0.05, **p<0.01, ***p<0.001.

**Supplementary Figure S2:**
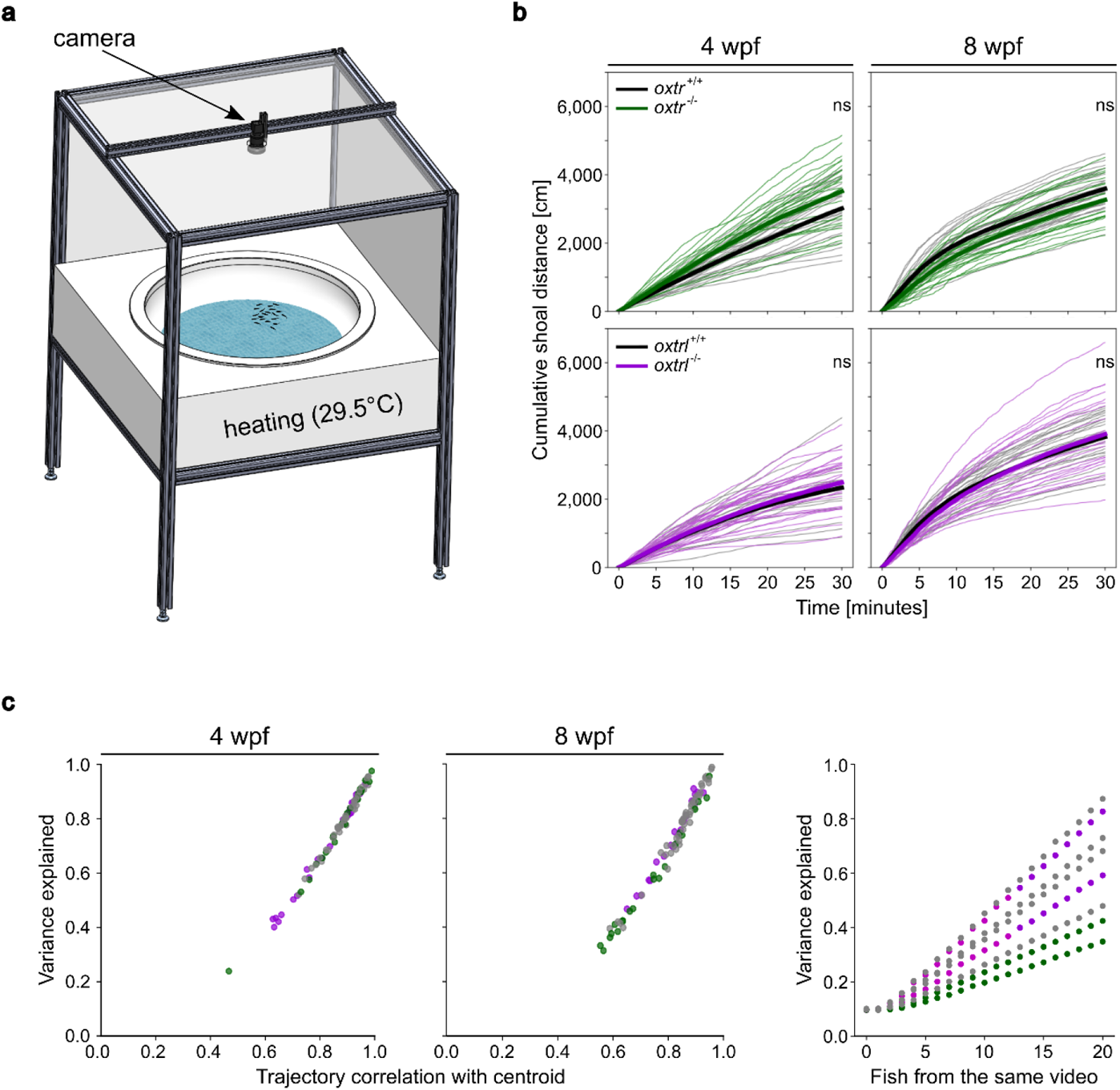
Shoaling setup details, cumulative shoal distance and variance explained as polarization parameter. **(a)** The behavioral chamber for the shoaling experiments was composed of a round tank, surrounded by 29.5°C warm water. White LEDs in the ceiling provided illumination for the camera positioned above. **(b)** Knocking-out one of the Oxytocin receptors did not lead to significant changes in the cumulative shoal distance at both 4 and 8 wpf. **(c)** The variance explained correlates with the trajectory correlation with centroid, making the variance explained a useful polarization parameter. **(d)** Adding more fish of the same video, while maintaining a group size of 20 with trajectories chosen from random videos, increased the variance explained – indicating its suitability as polarization parameter. Shown are two exemplar replicates per genotype group, color-coded in grey (*oxtr* ^+/+^ and *oxtrl* ^+/+^), green (*oxtr* ^-/-^) and purple (*oxtrl* ^-/-^). Number of replicates (n) can be found in Supplementary Table S1 and significance values in Supplementary Table S2. Significance is reported as *p<0.05, **p<0.01, ***p<0.001.

## Notes

### Competing Interest Statement

The authors have declared no competing interest.

