## Supplementary Table S1 for "Oxytocin receptors influence the development and maintenance of social behavior in zebrafish (*Danio rerio*)"

### Supplementary Table S1: Number of biological replicates.

**Social preference test** (Figures 1 and 2)

| rearing condition | social | | | | | | | | | | | |
|---|---|---|---|---|---|---|---|---|---|---|---|---|
| age group | 2 wpf | | | 3 wpf | | | 4 wpf | | | 8 wpf | | |
| genotype group | wildtype | *oxtr*$^{-/-}$ | *oxtrl*$^{-/-}$ | wildtype | *oxtr*$^{-/-}$ | *oxtrl*$^{-/-}$ | wildtype | *oxtr*$^{-/-}$ | *oxtrl*$^{-/-}$ | wildtype | *oxtr*$^{-/-}$ | *oxtrl*$^{-/-}$ |
| $n_{(inAnalysis)}$ | 15 | 15 | 18 | 15 | 18 | 18 | 16 | 12 | 15 | 13 | 16 | 11 |
| rearing condition | isolated | | | | | | | | | | | |
| age group | 2 wpf | | | 3 wpf | | | 4 wpf | | | 8 wpf | | |
| genotype group | wildtype | *oxtr*$^{-/-}$ | *oxtrl*$^{-/-}$ | wildtype | *oxtr*$^{-/-}$ | *oxtrl*$^{-/-}$ | wildtype | *oxtr*$^{-/-}$ | *oxtrl*$^{-/-}$ | wildtype | *oxtr*$^{-/-}$ | *oxtrl*$^{-/-}$ |
| $n_{(inAnalysis)}$ | 11 | 9 | 13 | 11 | 18 | 12 | 9 | 11 | 13 | 16 | 10 | 17 |

**Shoaling test** (Figures 3 and S2)

| age group | 4 wpf | | | | 8 wpf | | | |
|---|---|---|---|---|---|---|---|---|
| genotype group | *oxtr*$^{+/+}$ | *oxtr*$^{-/-}$ | *oxtrl*$^{+/+}$ | *oxtrl*$^{-/-}$ | *oxtr*$^{+/+}$ | *oxtr*$^{-/-}$ | *oxtrl*$^{+/+}$ | *oxtrl*$^{-/-}$ |
| $n_{(inAnalysis)}$ | 26 | 25 | 15 | 28 | 24 | 21 | 22 | 23 |

**qPCR** (Figure S1)

| group | *oxtrl* level | | *oxtr* level | | 4 wpf wildtype | | 8 wpf wildtype | |
|---|---|---|---|---|---|---|---|---|
| | *oxtr*$^{+/+}$ | *oxtr*$^{-/-}$ | *oxtrl*$^{+/+}$ | *oxtrl*$^{-/-}$ | social | isolated | social | isolated |
| $n_{(inAnalysis)}$ | 5 | 6 | 6 | 4 | 5 | 6 | 11 | 10 |
