## Supplementary Table S2 for "Oxytocin receptors influence the development and maintenance of social behavior in zebrafish (*Danio rerio*)"

**Supplementary Table S2: Significance values of statistical tests** (significance is reported as *p<0.05, **p<0.01 and ***p<0.001).

| | Kruskal-Wallis test | | | Wilcoxon rank-sum test | | |
|---|---|---|---|---|---|---|
| | groups | p value | significance | groups | p value | significance |
| Fig. 1b | All 2 wpf, socially reared: wildtype, *oxtr* $^{-/-}$, *oxtrl* $^{-/-}$ | $2.25 \times 10^{-1}$ | ns | All 2 wpf, socially reared: wildtype, *oxtr* $^{-/-}$ | $8.68 \times 10^{-1}$ | ns |
| | | | | All 2 wpf, socially reared: wildtype, *oxtrl* $^{-/-}$ | $1.47 \times 10^{-1}$ | ns |
| | | | | All 2 wpf, socially reared: *oxtr* $^{-/-}$, *oxtrl* $^{-/-}$ | $1.57 \times 10^{-1}$ | ns |
| | All 3 wpf, socially reared: wildtype, *oxtr* $^{-/-}$, *oxtrl* $^{-/-}$ | $2.25 \times 10^{-2}$ | * | All 3 wpf, socially reared: wildtype, *oxtr* $^{-/-}$ | $1.68 \times 10^{-2}$ | * |
| | | | | All 3 wpf, socially reared: wildtype, *oxtrl* $^{-/-}$ | $1.95 \times 10^{-2}$ | * |
| | | | | All 3 wpf, socially reared: *oxtr* $^{-/-}$, *oxtrl* $^{-/-}$ | $7.00 \times 10^{-1}$ | ns |
| | All 4 wpf, socially reared: wildtype, *oxtr* $^{-/-}$, *oxtrl* $^{-/-}$ | $3.61 \times 10^{-2}$ | * | All 4 wpf, socially reared: wildtype, *oxtr* $^{-/-}$ | $9.26 \times 10^{-1}$ | ns |
| | | | | All 4 wpf, socially reared: wildtype, *oxtrl* $^{-/-}$ | $2.41 \times 10^{-2}$ | * |
| | | | | All 4 wpf, socially reared: *oxtr* $^{-/-}$, *oxtrl* $^{-/-}$ | $3.56 \times 10^{-2}$ | * |
| | All 8 wpf, socially reared: wildtype, *oxtr* $^{-/-}$, *oxtrl* $^{-/-}$ | $2.87 \times 10^{-1}$ | ns | All 8 wpf, socially reared: wildtype, *oxtr* $^{-/-}$ | $8.95 \times 10^{-1}$ | ns |
| | | | | All 8 wpf, socially reared: wildtype, *oxtrl* $^{-/-}$ | $2.83 \times 10^{-1}$ | ns |
| | | | | All 8 wpf, socially reared: *oxtr* $^{-/-}$, *oxtrl* $^{-/-}$ | $1.14 \times 10^{-1}$ | ns |
| Fig. 1c | All wildtype, socially reared: 2, 3, 4 and 8 wpf | $9.31 \times 10^{-1}$ | ns | All wildtype, socially reared: 2 wpf, 3 wpf | $8.03 \times 10^{-1}$ | ns |
| | | | | All wildtype, socially reared: 2 wpf, 4 wpf | $5.93 \times 10^{-1}$ | ns |

| | | | | | | |
|---|---|---|---|---|---|---|
| Fig. 1c | All wildtype, socially reared: 2, 3, 4 and 8 wpf | $9.31 \times 10^{-1}$ | ns | All wildtype, socially reared: 2 wpf, 8 wpf | $7.47 \times 10^{-1}$ | ns |
| | | | | All wildtype, socially reared: 3 wpf, 4 wpf | $7.07 \times 10^{-1}$ | ns |
| | | | | All wildtype, socially reared: 3 wpf, 8 wpf | $6.95 \times 10^{-1}$ | ns |
| | | | | All wildtype, socially reared: 4 wpf, 8 wpf | $7.59 \times 10^{-1}$ | ns |
| | All *oxtr* $^{-/-}$, socially reared: 2, 3, 4 and 8 wpf | $1.65 \times 10^{-2}$ | * | All *oxtr* $^{-/-}$, socially reared: 2 wpf, 3 wpf | $6.10 \times 10^{-3}$ | ** |
| | | | | All *oxtr* $^{-/-}$, socially reared: 2 wpf, 4 wpf | $4.07 \times 10^{-1}$ | ns |
| | | | | All *oxtr* $^{-/-}$, socially reared: 2 wpf, 8 wpf | $6.07 \times 10^{-1}$ | ns |
| | | | | All *oxtr* $^{-/-}$, socially reared: 3 wpf, 4 wpf | $3.70 \times 10^{-2}$ | * |
| | | | | All *oxtr* $^{-/-}$, socially reared: 3 wpf, 8 wpf | $1.65 \times 10^{-2}$ | * |
| | | | | All *oxtr* $^{-/-}$, socially reared: 4 wpf, 8 wpf | $6.42 \times 10^{-1}$ | ns |
| | All *oxtrl* $^{-/-}$, socially reared: 2, 3, 4 and 8 wpf | $4.09 \times 10^{-5}$ | *** | All *oxtrl* $^{-/-}$, socially reared: 2 wpf, 3 wpf | $3.49 \times 10^{-5}$ | *** |
| | | | | All *oxtrl* $^{-/-}$, socially reared: 2 wpf, 4 wpf | $1.18 \times 10^{-4}$ | *** |
| | | | | All *oxtrl* $^{-/-}$, socially reared: 2 wpf, 8 wpf | $3.60 \times 10^{-3}$ | ** |
| | | | | All *oxtrl* $^{-/-}$, socially reared: 3 wpf, 4 wpf | $9.86 \times 10^{-1}$ | ns |
| | | | | All *oxtrl* $^{-/-}$, socially reared: 3 wpf, 8 wpf | $7.01 \times 10^{-1}$ | ns |
| | | | | All *oxtrl* $^{-/-}$, socially reared: 4 wpf, 8 wpf | $9.38 \times 10^{-1}$ | ns |

| | | | | | | |
|---|---|---|---|---|---|---|
| | All 2 wpf, isolated reared: wildtype, *oxtr* $^{-/-}$, *oxtrl* $^{-/-}$ | $1.85 \times 10^{-1}$ | ns | All 2 wpf, isolated reared: wildtype, *oxtr* $^{-/-}$ | $1.11 \times 10^{-1}$ | ns |
| | | | | All 2 wpf, isolated reared: wildtype, *oxtrl* $^{-/-}$ | $1.48 \times 10^{-1}$ | ns |
| | | | | All 2 wpf, isolated reared: *oxtr* $^{-/-}$, *oxtrl* $^{-/-}$ | $6.89 \times 10^{-1}$ | ns |
| | All 3 wpf, isolated reared: wildtype, *oxtr* $^{-/-}$, *oxtrl* $^{-/-}$ | $3.51 \times 10^{-1}$ | ns | All 3 wpf, isolated reared: wildtype, *oxtr* $^{-/-}$ | $1.44 \times 10^{-1}$ | ns |
| | | | | All 3 wpf, isolated reared: wildtype, *oxtrl* $^{-/-}$ | $4.06 \times 10^{-1}$ | ns |
| | | | | All 3 wpf, isolated reared: *oxtr* $^{-/-}$, *oxtrl* $^{-/-}$ | $7.19 \times 10^{-1}$ | ns |
| Fig. 2b | All 4 wpf, isolated reared: wildtype, *oxtr* $^{-/-}$, *oxtrl* $^{-/-}$ | $1.80 \times 10^{-3}$ | ** | All 4 wpf, isolated reared: wildtype, *oxtr* $^{-/-}$ | $3.34 \times 10^{-2}$ | * |
| | | | | All 4 wpf, isolated reared: wildtype, *oxtrl* $^{-/-}$ | $8.41 \times 10^{-4}$ | *** |
| | | | | All 4 wpf, isolated reared: *oxtr* $^{-/-}$, *oxtrl* $^{-/-}$ | $9.29 \times 10^{-2}$ | ns |
| | All 8 wpf, isolated reared: wildtype, *oxtr* $^{-/-}$, *oxtrl* $^{-/-}$ | $4.82 \times 10^{-1}$ | ns | All 8 wpf, isolated reared: wildtype, *oxtr* $^{-/-}$ | $6.54 \times 10^{-1}$ | ns |
| | | | | All 8 wpf, isolated reared: wildtype, *oxtrl* $^{-/-}$ | $2.14 \times 10^{-1}$ | ns |
| | | | | All 8 wpf, isolated reared: *oxtr* $^{-/-}$, *oxtrl* $^{-/-}$ | $7.07 \times 10^{-1}$ | ns |
| Fig. 2c | | | | All 2 wpf wildtype: social, isolated | $2.54 \times 10^{-1}$ | ns |
| | | | | All 3 wpf wildtype: social, isolated | $9.59 \times 10^{-1}$ | ns |
| | | | | All 4 wpf wildtype: social, isolated | $2.03 \times 10^{-1}$ | ns |
| | | | | All 8 wpf wildtype: social, isolated | $4.10 \times 10^{-3}$ | ** |

| | | | | |
|---|---|---|---|---|
| Fig. 2c | | All 2 wpf *oxtr* $^{-/-}$: social, isolated | $4.09 \times 10^{-1}$ | ns |
| | | All 3 wpf *oxtr* $^{-/-}$: social, isolated | $3.45 \times 10^{-1}$ | ns |
| | | All 4 wpf *oxtr* $^{-/-}$: social, isolated | $2.82 \times 10^{-1}$ | ns |
| | | All 8 wpf *oxtr* $^{-/-}$: social, isolated | $6.90 \times 10^{-2}$ | ns |
| | | All 2 wpf *oxtrl* $^{-/-}$: social, isolated | $6.82 \times 10^{-2}$ | ns |
| | | All 3 wpf *oxtrl* $^{-/-}$: social, isolated | $3.50 \times 10^{-1}$ | ns |
| | | All 4 wpf *oxtrl* $^{-/-}$: social, isolated | $8.94 \times 10^{-5}$ | *** |
| | | All 8 wpf *oxtrl* $^{-/-}$: social, isolated | $7.30 \times 10^{-3}$ | ** |
| Fig. 3a | | All 4 wpf, socially reared: *oxtr* $^{+/+}$, *oxtr* $^{-/-}$ | $4.74 \times 10^{-1}$ | ns |
| | | All 4 wpf, socially reared: *oxtrl* $^{+/+}$, *oxtrl* $^{-/-}$ | $2.11 \times 10^{-1}$ | ns |
| | | All 4 wpf, socially reared: *oxtr* $^{+/+}$, *oxtrl* $^{+/+}$ | $1.04 \times 10^{-1}$ | ns |
| | | All 8 wpf, socially reared: *oxtr* $^{+/+}$, *oxtr* $^{-/-}$ | $5.48 \times 10^{-4}$ | *** |
| | | All 8 wpf, socially reared: *oxtrl* $^{+/+}$, *oxtrl* $^{-/-}$ | $2.75 \times 10^{-2}$ | * |
| | | All 8 wpf, socially reared: *oxtr* $^{+/+}$, *oxtrl* $^{+/+}$ | $5.30 \times 10^{-2}$ | ns |
| Fig. 3b | | All 4 wpf, socially reared: *oxtr* $^{+/+}$, *oxtr* $^{-/-}$ | $3.85 \times 10^{-1}$ | ns |

| | | | | |
|---|---|---|---|---|
| Fig. 3b | | All 4 wpf, socially reared: *oxtrl* $^{+/+}$, *oxtrl* $^{-/-}$ | 8.64 x $10^{-2}$ | ns |
| | | All 4 wpf, socially reared: *oxtr* $^{+/+}$, *oxtrl* $^{+/+}$ | 4.09 x $10^{-1}$ | ns |
| | | All 8 wpf, socially reared: *oxtr* $^{+/+}$, *oxtr* $^{-/-}$ | 4.59 x $10^{-3}$ | ** |
| | | All 8 wpf, socially reared: *oxtrl* $^{+/+}$, *oxtrl* $^{-/-}$ | 1.34 x $10^{-2}$ | * |
| | | All 8 wpf, socially reared: *oxtr* $^{+/+}$, *oxtrl* $^{+/+}$ | 8.47 x $10^{-2}$ | ns |
| Fig. 3c | | All 4 wpf, socially reared: *oxtr* $^{+/+}$, *oxtr* $^{-/-}$ | 4.74 x $10^{-1}$ | ns |
| | | All 4 wpf, socially reared: *oxtrl* $^{+/+}$, *oxtrl* $^{-/-}$ | 7.86 x $10^{-2}$ | ns |
| | | All 4 wpf, socially reared: *oxtr* $^{+/+}$, *oxtrl* $^{+/+}$ | 8.58 x $10^{-2}$ | ns |
| | | All 8 wpf, socially reared: *oxtr* $^{+/+}$, *oxtr* $^{-/-}$ | 8.19 x $10^{-3}$ | ** |
| | | All 8 wpf, socially reared: *oxtrl* $^{+/+}$, *oxtrl* $^{-/-}$ | 1.27 x $10^{-2}$ | * |
| | | All 8 wpf, socially reared: *oxtr* $^{+/+}$, *oxtrl* $^{+/+}$ | 5.30 x $10^{-2}$ | ns |
| Fig. 3d | | All 4 wpf, socially reared: *oxtr* $^{+/+}$, *oxtr* $^{-/-}$ | 4.44 x $10^{-1}$ | ns |
| | | All 4 wpf, socially reared: *oxtrl* $^{+/+}$, *oxtrl* $^{-/-}$ | 5.28 x $10^{-2}$ | ns |
| | | All 4 wpf, socially reared: *oxtr* $^{+/+}$, *oxtrl* $^{+/+}$ | 1.90 x $10^{-1}$ | ns |
| | | All 8 wpf, socially reared: *oxtr* $^{+/+}$, *oxtr* $^{-/-}$ | 5.24 x $10^{-3}$ | ** |

| | | | | |
|---|---|---|---|---|
| Fig. 3d | | All 8 wpf, socially reared: *oxtrl* $^{+/+}$, *oxtrl* $^{-/-}$ | $1.79 \times 10^{-2}$ | * |
| | | All 8 wpf, socially reared: *oxtr* $^{+/+}$, *oxtrl* $^{+/+}$ | $2.33 \times 10^{-2}$ | * |
| Fig. S1b | | All 8 wpf, socially reared: *oxtr* $^{+/+}$, *oxtr* $^{-/-}$ | $9.31 \times 10^{-1}$ | ns |
| | | All 8 wpf, socially reared: *oxtrl* $^{+/+}$, *oxtrl* $^{-/-}$ | $6.10 \times 10^{-1}$ | ns |
| Fig. S1c | | All 4 wpf wildtype: social, isolated | $3.29 \times 10^{-1}$ | ns |
| | | All 8 wpf wildtype: social, isolated | $6.70 \times 10^{-3}$ | ** |
| | | All wildtype, socially reared: 4 wpf, 8 wpf | $4.58 \times 10^{-4}$ | *** |
| | | All wildtype, isolated reared: 4 wpf, 8 wpf | $3.13 \times 10^{-1}$ | ns |
| Fig. S2b | | All 4 wpf, socially reared: *oxtr* $^{+/+}$, *oxtr* $^{-/-}$ | $5.34 \times 10^{-2}$ | ns |
| | | All 4 wpf, socially reared: *oxtrl* $^{+/+}$, *oxtrl* $^{-/-}$ | $4.52 \times 10^{-1}$ | ns |
| | | All 4 wpf, socially reared: *oxtr* $^{+/+}$, *oxtrl* $^{+/+}$ | $1.79 \times 10^{-2}$ | * |
| | | All 8 wpf, socially reared: *oxtr* $^{+/+}$, *oxtr* $^{-/-}$ | $6.05 \times 10^{-2}$ | ns |
| | | All 8 wpf, socially reared: *oxtrl* $^{+/+}$, *oxtrl* $^{-/-}$ | $9.37 \times 10^{-1}$ | ns |
| | | All 8 wpf, socially reared: *oxtr* $^{+/+}$, *oxtrl* $^{+/+}$ | $1.50 \times 10^{-1}$ | ns |
